# Novel Engineered AAV Variants Demonstrate Superior Blood-Brain Barrier Penetration and Safety in Non-Human Primates

**DOI:** 10.64898/2026.03.29.713052

**Authors:** Zhenhua Wang, Hao Li, Xianggang Xu, Zhen Sun, Rui He, Linlin Zhang, Mengmeng Yu, Shengwen Wang, Can Hu, Lei Liu, Lele Ren, Yanling Xu, Tongqian Xiao, Daoyuan Li, Bo Sun, Youguang Luo, Zhenming An

**Author notes:** Correspondence: Youguang Luo,; Bo Sun,; Zhenming An. These authors contributed equally (Zhenhua Wang, Hao Li, Xianggang Xu, Zhen Sun).

## Abstract

Systemic delivery of adeno-associated virus (AAV) for gene therapy of central nervous system (CNS) disorders is limited by inefficient blood-brain barrier (BBB) penetration and dose-limiting toxicity in peripheral organs, notably the liver and dorsal root ganglia (DRG)^1-5^. Here, we report the development of novel AAV variants via a proprietary capsid engineering platform (REACH). In non-human primates (NHPs), intravenous administration of lead variants resulted in transgene expression levels in the brain that were 600-2000 fold higher than AAV9 at the RNA level, concomitant with a 10-50 fold reduction in liver tropism and minimal off-target exposure in the heart and DRG. These engineered capsids achieve unprecedented, pan-CNS transduction with a markedly improved safety profile, representing a transformative platform for treating a broad spectrum of neurological diseases.

## Main

### 1. Proprietary REACH AAV Delivery Platform

Gene therapy holds immense promise for treating monogenic and complex disorders of the CNS. However, the BBB presents a formidable obstacle to the delivery of therapeutic agents. While some engineered AAV capsids show enhanced CNS transduction in rodents, their performance and safety in higher-order species, critical for clinical translation, remain inadequately characterized^6,7^. A major challenge is the pervasive trade-off between enhanced brain delivery and increased off-target exposure, particularly in the liver and DRG, which raises significant safety concerns.

To overcome these limitations, we employed a combined strategy of rational design and directed evolution (REACH platform) to generate a diverse library of AAV-based capsids. And then a multi-tiered screening system, including in vitro screening, in vivo screening and leads confirmation, was used to identify the top candidates (Fig. 1).

**Fig. 1.**
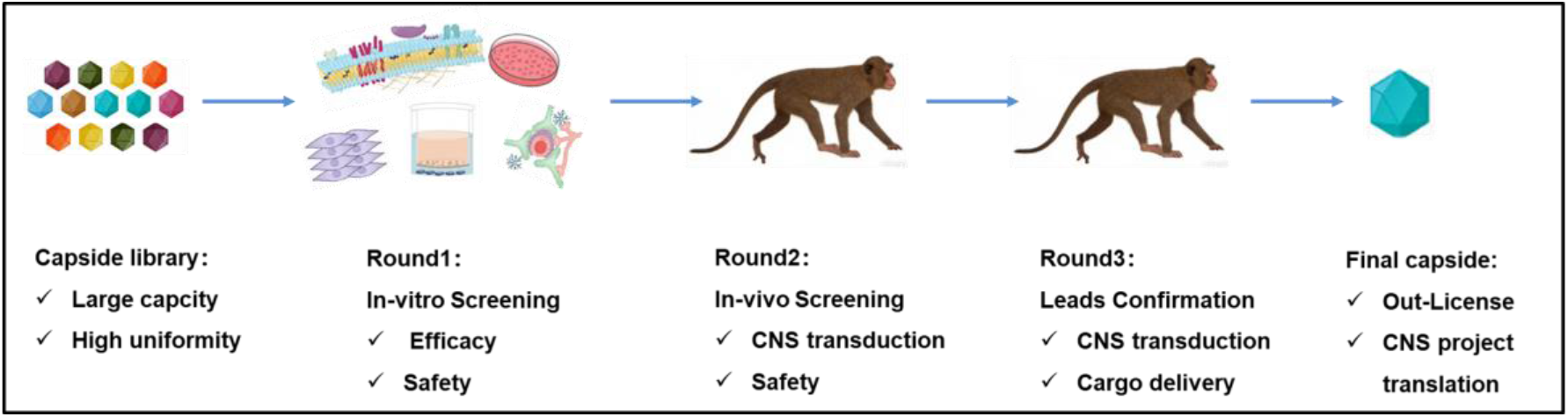
Schematic of the engineered AAV capsid screening pipeline based on REACH platform. A high-capacity, highly uniform AAV library was constructed de novo. The primary screening was performed through in vitro validation, yielding a lead library for next screening. This enriched library was then administered intravenously to non-human primates (NHPs) for a secondary in vivo screening round, leading to the identification of a series of highly efficient, blood-brain barrier-penetrant AAV variants targeting the central nervous system. The top-performing variants were further characterized individually with a therapeutic cargo in NHPs to get more comprehensive efficacy and safety profile, establishing a foundation for subsequent clinical translation.

### 2. Engineered AAV Variants Enable Efficient Blood-Brain Barrier Crossing

Quantification of AAV genomes (DNA) and transgene mRNA (RNA) in the NHP brain samples revealed a dramatic enhancement in delivery efficiency with next-generation sequencing (NGS) analysis. The 5 lead variants (C10, C28, C11, C29, C24) exhibited a 50-130 fold increase in DNA distribution and a staggering 600-2000 fold increase in RNA expression compared to AAV9, significantly outperforming the published positive controls (Fig. 2).

**Fig. 2.**
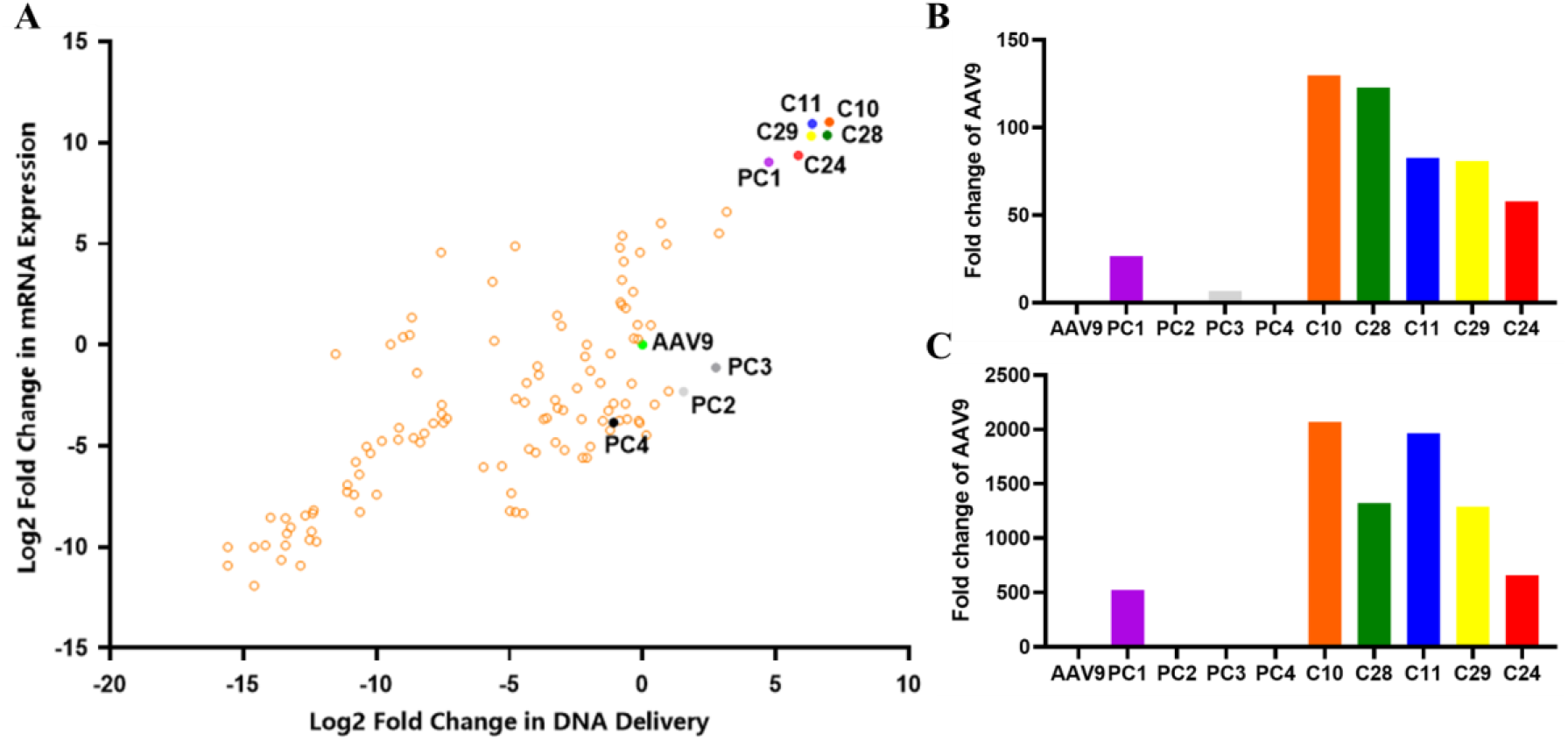
Engineered AAV variants enable efficient blood-brain barrier crossing. (A) Distribution of AAV variants in the NHP brain. In NHP studies, AAV variants were intravenous injected and then the DNA and RNA level of each AAV variant from brain samples are analyzed with NGS sequencing.

### 3. Engineered AAV Variants Achieve Widespread Brain Distribution

NGS analysis of AAV genome and transgene-derived RNA demonstrated that the lead variants mediate broad, pan-CNS transduction. High-level expression was consistently observed across all major brain regions examined, including the cortex, cerebellum, brainstem, hippocampus and deep brain, with peak levels in the cortex (Fig. 3). This uniform distribution highlights the potential of these capsids as universal vectors for global CNS gene delivery.

**Fig. 3.**
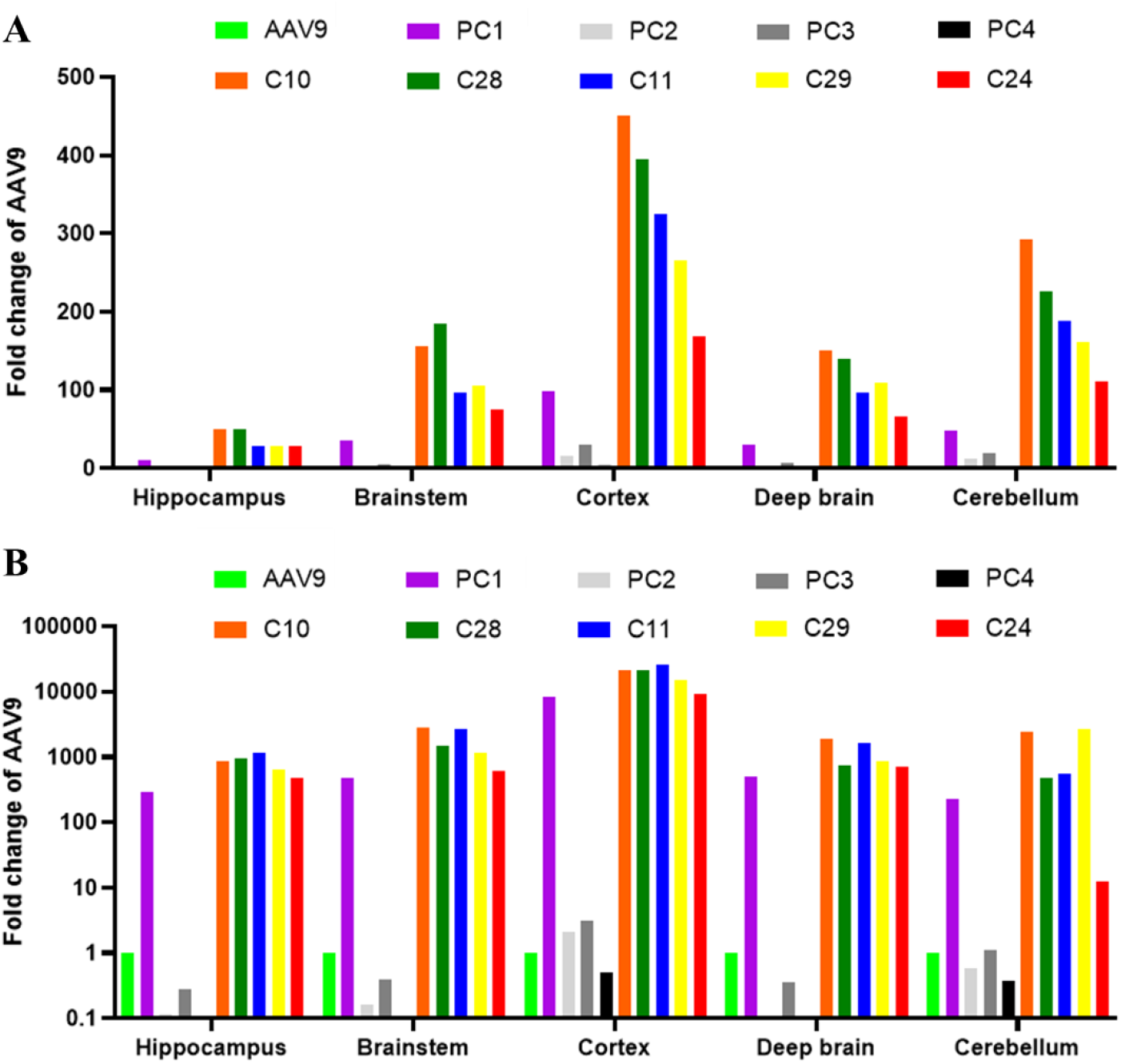
Engineered AAV variants demonstrate widespread distribution and enhanced delivery across NHP brain regions. (A) DNA data of AAV distribution in the NHP brain regions. In NHP studies, AAV variants were intravenous injected and then the DNA level of each AAV variant from brain region samples are analyzed with NGS sequencing. Data are presented as fold change relative to AAV9. (B) RNA data of AAV distribution in the NHP brain regions. In NHP studies, AAV variants were intravenous injected and then the RNA level of each AAV variant from brain region

### 4. Engineered AAV Variants Show a Favorable Safety Profile

Crucially, this exceptional CNS transduction is not achieved at the expense of systemic safety. Biodistribution analysis demonstrated a highly favorable profile for the lead variants. Hepatic exposure was reduced by 10-50 fold at the DNA level compared to AAV9 (Fig. 4A). Furthermore, off-target exposure in critical tissues, such as the DRG and heart, was minimal, in stark contrast to the signals observed for AAV9 and most of the positive controls (Fig. 4B, C).

**Fig. 4.**
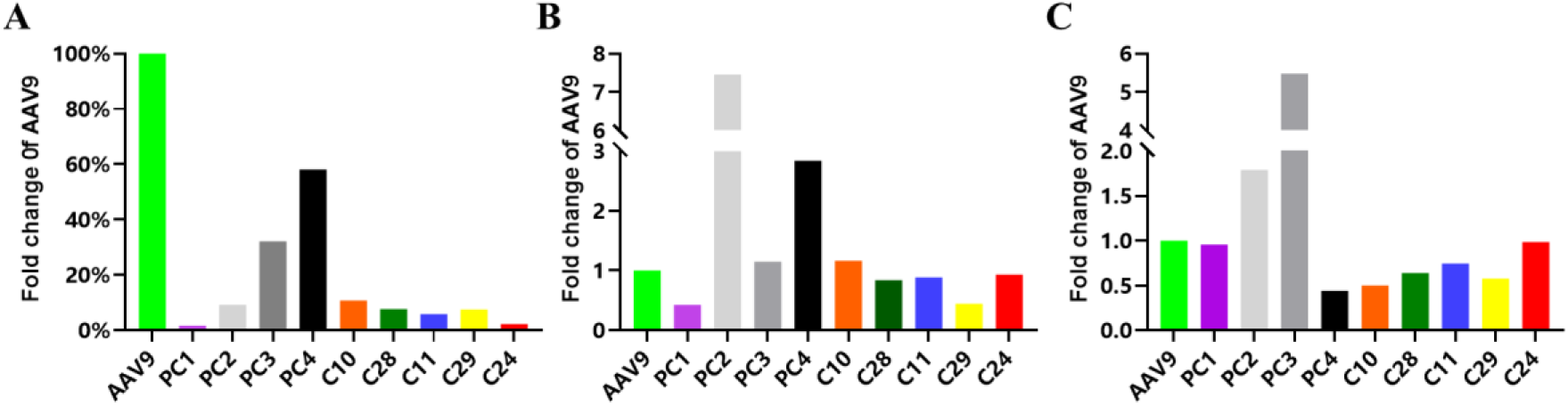
Engineered AAV variants show a favorable safety profile. . (A) Fold change in AAV distribution in the liver. (B) Fold change in AAV distribution in the DRG. (C) Fold change in AAV distribution in the heart. NHPs were intravenously injected with indicated AAV samples. Tissues of interest, such as liver, DRG, and heart, were collected and detected for viral genome copy numbers with NGS sequencing, and the fold change for each variant relative to AAV9 was calculated. PC indicates positive control.

## Discussion/Overall summary

We have developed a new class of AAV-based capsids that simultaneously break the efficiency ceiling and raise the safety floor for systemic CNS gene therapy. The magnitude of improvement, up to 2000 fold in CNS transgene expression, represents a qualitative leap over existing technologies^8-12^. More importantly, this efficacy is coupled with a significant de-targeting of the liver and avoidance of DRG and heart, effectively widening the therapeutic window.

The NHP data presented here are of direct translational relevance, addressing the primary efficacy and safety bottlenecks that have hindered the clinical advancement of systemic CNS gene therapies. The lead variants emerged from a platform designed to bridge in vitro human cell models and in vivo primate physiology, which may explain their superior performance.

These engineered capsids constitute a versatile platform capable of delivering diverse genetic cargoes. Their ability to achieve efficient, safe, and pan-CNS transduction following a simple intravenous injection could accelerate the clinical translation of therapies for a wide range of neurological conditions.

### Here is the take-home message

- Five novel AAV variants with significantly higher BBB-crossing efficiency were found with Qilu’s proprietary REACH platform.
- These lead capsids show superior CNS transduction in NHPs, with 50-130 fold enhanced DNA distribution and 600-2000 fold increased RNA expression versus AAV9.
- These lead capsids exhibit significantly improved safety, achieving 10-50 fold liver de-targeting while maintaining or reducing exposure in peripheral tissues, such as the heart and DRG.
- These best-in-class BBB-penetrating vectors represent a transformative platform for CNS gene therapy, with robust NHP data supporting immediate partnership opportunities for clinical translation and out-licensing.

## Online content

Any methods, additional references, extended data, supplementary information, acknowledgements, author contributions and competing interests are available online.

## Methods

### AAV sample preparation for NHP study

All AAV variants were produced using the standard triple-plasmid transfection method in suspension HEK293 cells.

- **Cell Culture and Transfection:** HEK293 cells were passaged when density reached
- ≥3×10^6^ cells/mL. For transfection, cells were seeded at 2×10^6^ cells/mL. A mixture of three plasmids—the adenoviral helper plasmid, the AAV rep/cap plasmid, and the transgene plasmid encoding enhanced green fluorescent protein with a barcode tag (EGFP-Barcode) —was combined at a 1:1:1 molar ratio. The DNA was mixed with polyethylenimine (PEI) at a PEI:DNA=2:1 ratio (v/w).
- **Harvest and Lysis:** Three days post-transfection, cells were lysed by adding a lysis buffer containing Tris, MgCl_2_, and Tween-20, along with Benzonase (50 U/mL) to digest unpackaged nucleic acids. The lysate was incubated at 37°C for 3 hours and then clarified by centrifugation.
- **Purification:** The clarified harvest was filtered and subjected to affinity chromatography using an AAVX resin. The eluted virus sample was further purified by iodixanol density gradient ultracentrifugation. The final viral band was collected, buffer-exchanged, and concentrated using tangential flow filtration (TFF).
- **AAV Virus Release Test:** The titer of AAV sample was detected by quantitative real-time polymerase chain reaction (qPCR). The particle size distribution was analyzed by dynamic light scattering (DLS). The purity were analyzed by SDS-PAGE and SEC. Endotoxin level, and sterility were also detected according to the SOPs.

### AAV BBB-penetrating efficacy and safety profile evaluation in NHP

In NHP studies, indicated AAV variants were intravenous injected and then the DNA and RNA level of each AAV variant from samples of interest were analyzed with NGS sequencing.

- **NHP pre-screening:** The study was conducted in cynomolgus macaques (Macaca fascicularis). Prior to dosing, animals were screened for pre-existing neutralizing antibodies (Nabs) against AAV, and monkeys with lower Nab titers were selected.
- **Dosing and Tissue Processing:** Each animal received a single intravenous infusion of the pooled AAV library. 28 days post-dosing, animals were perfused and necropsied. Tissues, including multiple brain regions (hippocampus, cortex, deep brain, cerebellum, and brainstem), liver, and DRG, were collected, snap-frozen, and stored at -80°C for further study.
- **Biodistribution Analysis via NGS:** Genomic DNA or RNA sample was extracted from all tissues of interest. The region containing the unique barcode was amplified by PCR from each sample, and the resulting amplicons were subjected to NGS analysis. The relative abundance of each barcode (corresponding to each AAV variant) in every tissue was calculated from the NGS read counts.

## Data availability

The data that support the findings of this study are available from the corresponding author upon reasonable request.

## Acknowledgements

This study was funded by Qilu Pharmaceutical Co., LTD. The authors thank the Institute of Biopharmaceuticals team, pre-clinical team et al. for their contribution in designing/performing experiments and data analysis.

## Author contributions

Conceptualization: ZW, YL, ZA; Methodology: ZW, YL, LZ, YX, TX, HL, DL; Experimentation: ZW, XX, ZS, HL, RH, MY, SW, CH, LL, LR; Resources: ZA, BS, YL; Formal Analysis: ZW, RH, ZS, XX, LZ; Investigation: ZW; Writing-Original Draft: ZW, HL, XX, ZS, RH; Writing-Review & Editing: YL, ZA.

